# Genomic surveillance unfolds the dynamics of SARS-CoV-2 transmission and divergence in Bangladesh over the past two years

**DOI:** 10.1101/2022.04.13.488264

**Authors:** Tushar Ahmed Shishir, Taslimun Jannat, Iftekhar Bin Naser

**Author notes:** Address correspondence to: Iftekhar Bin Naser.

## Abstract

The highly pathogenic virus SARS-CoV-2 has shattered the healthcare system of the world causing the COVID-19 pandemic since first detected in Wuhan, China. Therefore, scrutinizing the genome structure and tracing the transmission of the virus has gained enormous interest in designing appropriate intervention strategies to control the pandemic. In this report, we examined 4622 sequences from Bangladesh and found that they belonged to thirty-five major PANGO lineages, while Delta alone accounted for 39%, and 78% were from just four primary lineages. Our research has also shown Dhaka to be the hub of viral transmission and observed the virus spreading back and forth across the country at different times by building a transmission network. The analysis resulted in 7659 unique mutations, with an average of 24.61 missense mutations per sequence. Moreover, our analysis of genetic diversity and mutation patterns revealed that eight genes were under negative selection pressure to purify deleterious mutations, while three genes were under positive selection pressure.

**Importance:** With 29,122 deaths, 1.95 million infections and a shattered healthcare system from SARS-CoV-2 in Bangladesh, the only way to avoid further complications is to break the transmission network of the virus. Therefore, it is vital to shedding light on the transmission, divergence, mutations, and emergence of new variants using genomic data analyses and surveillance. Here, we present the geographic and temporal distribution of different SARS-CoV-2 variants throughout Bangladesh over the past two years, and their current prevalence. Further, we have developed a transmission network of viral spreads, which in turn will help take intervention measures. Then we analyzed all the mutations that occurred and their effect on evolution as well as the currently present mutations that could trigger a new variant of concern. In short, together with an ongoing genomic surveillance program, these data will help to better understand SARS-CoV-2, its evolution, and pandemic characteristics in Bangladesh.

## Introduction

The Coronavirus disease (COVID-19) pandemic was initially reported as an unknown respiratory illness towards the end of 2019. However, it eventually became evident that a novel SARS-like coronavirus was causing the infections and the virus was termed severe acute respiratory syndrome coronavirus 2 (SARS-CoV-2) (1). Originating in Wuhan, China, SARS-CoV-2 has spread across 220 countries and territories, infecting 488.75 million and causing the death of 6.17 million people till 31st March 2022, resulting in a global economic crisis, which is the third zoonotic virus after MERS-CoV and SARS-CoV in 2012 and 2002 respectively (2, 3). The novel virus belonging to the Betacoronavirus genus and Coronaviridae family is a positive-sense, single-stranded ∼30 kb long RNA virus. Its genome contains 38% GC content (4), prefers pyrimidine-rich codons over purines (5) and is organized into 11 open reading frames expressing 12 proteins, including two polypeptides, four structural proteins and other accessory proteins (6). Phylogenetically, the virus shares 96% identity with the strain BatCoV RaTG13 of Rhinolophus affinis, and genome sequences along with epidemiological data suggest that SARS-CoV-2 is primarily transmitted from bats to humans (3, 4, 7). A complete genome sequence of the virus was deposited in GenBank on 5th January (NC_045512.2) (8), followed by the submission of 9.74 million complete sequences to GISAID by 25th March 2022 (9).

According to recent data from Worldometer, the most infected countries are the USA, India, and Brazil, with more than 29, 43, and 33 million cases of infection and thousands of deaths (2, 10). Since the first case was confirmed in Bangladesh on 8th March 2020, there have been 1.95 million positives and 29,122 deaths reported until 31st March 2022 (11). Having such a large population makes Bangladesh more vulnerable to viral transmission, and it is labelled as the second-most infected nation in the South Asian region (10), despite the government imposing lockdowns, social distancing rules and mask mandates to control the situation. Therefore, it is crucial to shed light on the transmission and evolution of the virus inside the country to reduce the fatality where genomic data analyses and surveillance comes into play, which can deliver immense information. Child Health Research Foundation published the first SARS-CoV-2 genome sequence from Bangladesh on 12th May 2020 (12), followed by 5146 further sequences until 25th March 2022 (9).

To date, Bangladesh has been affected by three waves of COVID-19 with variants of concerns (VOC), including Alpha, Beta, Delta, and Omicron (9). VOC is the name given to a variant of the SARS-CoV-2 virus that has mutations in the spike protein receptor-binding domain, increasing binding affinity within the RBD-hACE2 complex and increasing viral transmission (13, 14). Consequently, the mutations are essential for studying since they alter the antigenic potentials of the epitopes and consequently affect pathogenicity, infectivity, transmissibility, and evading host immunity. SARS-CoV-2 encodes an exoribonuclease that proofreads the errors during viral RNA synthesis; therefore, it has a lower mutation rate than other RNA viruses, which aids in enhancing its ability to adapt to their environment (15, 16). Nevertheless, the virus is accumulating mutations across its genome, leading to the emergence of different variants over time. These mutations are not evenly distributed; for example, some genes are more prone to mutations than others are, and cytosine to uracil substitution is more common in SARS-CoV-2, reforming the transition/transversion ratio, which is negatively correlated with evolutionary time (17). Additionally, a variable vaccination rate among the countries increases the risk of SARS-CoV-2 mutating into a strain that is resistant to current vaccines and therapies. Consequently, it is essential to continue investigating the mutations of SARS-CoV-2 in order to develop further effective vaccines and therapies, improve pandemic response, and reduce the impact of the pandemic on healthcare and clinical processes.

To the best of our knowledge, most previous studies in Bangladesh addressed lineages distribution, source determination, and potential mutations with only a few sequences from the early phase of the outbreak (18, 19). Therefore, in this work, we comprehensively analyzed 4622 whole-genome sequences of SARS-CoV-2 isolated from Bangladesh until 25th March 2022 to understand the distribution of variants and mutation accumulation trends over the year. We have thoroughly studied the temporal and geographical distribution of different lineages inside Bangladesh and built the transmission network to trace their back and forth circulation. To better understand the evolutionary dynamics of SARS-CoV-2 in Bangladesh over the last two years, we examined the genetic diversity among strains, gene-wise mutation distribution, and selection pressures.

## Results

### SARS-CoV-2 lineage dynamics

To understand the diversity and transmission of the virus, we have confined and analyzed sequences from all administrative divisions of Bangladesh. There were 5146 sequences submitted in GISAID till 25th March 2022, but many of them were incomplete and lacked quality. Therefore, we filtered the sequences based on their completeness, coverage, and gaps, resulting in 4892 sequences for downstream analysis (Supplementary file 1). However, when we examined PANGO Lineages, we observed that there were 93 lineages, many of which carried extremely low numbers of the sequences. Hence, we further filtered the sequences and kept only the lineages containing at least ten sequences, resulting in 4622 sequences from 35 lineages. Overall, in the beginning, the country had strains that belonged to the fewest number of PANGO lineages, but this has changed over time (Supplementary file 2). Selected sequences belonging to thirty-five different PANGO lineages provided us with invaluable insight regarding patterns of pandemic and viral spread (Supplementary file 2). As an example, 78% of the sequences were grouped into four lineages, where Delta (B.1.617.2) and its three major sub-lineages (AY.X) combined made up the highest 39% of the total sequences, while 20 out of thirty-five lineages held only 9% of sequences even after we filtered out the lineages with very few sequences. The top ten most prevalent lineages were found to be B.1.617.2 (23.34%), B.1.1.25 (20.68%), BA.2 (8.57%), B.1.351.3 (7.27%), B.1.1 (2.40%), B.1.1.7 (1.86%), B.1 (1.80%), B.1.351 (1.02%), B.1.36.16 (0.93%) and B.1.1.318 (0.74%).

### Temporal distribution of major lineages

We found that Bangladesh was afflicted by a large number of viruses from 35 different lineages, with the highest diversification occurring between July and September of 2021 with sequences from 20 to 23 lineages (supplementary file 2). The early phase of the pandemic in Bangladesh was started by the introduction of lineage B.1 in March 2020. Multiple occurrences of the introduction of COVID-19 from different countries have previously been reported; for instance, Dhaka was first exposed to COVID-19 with strains from the United Kingdom, while Chattogram was exposed to strains from Saudi Arabia (18). The early phase of the pandemic was generally dominated by imported strains from outside countries, but as the pandemic progressed, mutations changed dynamics and the linage B.1.1.25 took over, with B.1 gradually declining (Fig 1A). B.1.1.25 was the highest prevalent strain until January 2021. Later, the Beta variant (B.1.351) was reported in November 2020, followed by the Alpha variant (B.1.1.7) in December 2020. The B.1.1.7 lineage started talking over the B.1.1.25 lineage following its introduction. This linage was the most frequently detected variant in February 2021, while Beta variants were very less numerous. Despite this, a sub-lineage of Beta variants (B.1.351.3) emerged and outnumbered the Alpha variant in March 2021 (Supplementary file 2). However, the dominance of B.1.351.3 did not last long due to the introduction of the deadly delta variant (B.1.617.2).

**Fig 1:**
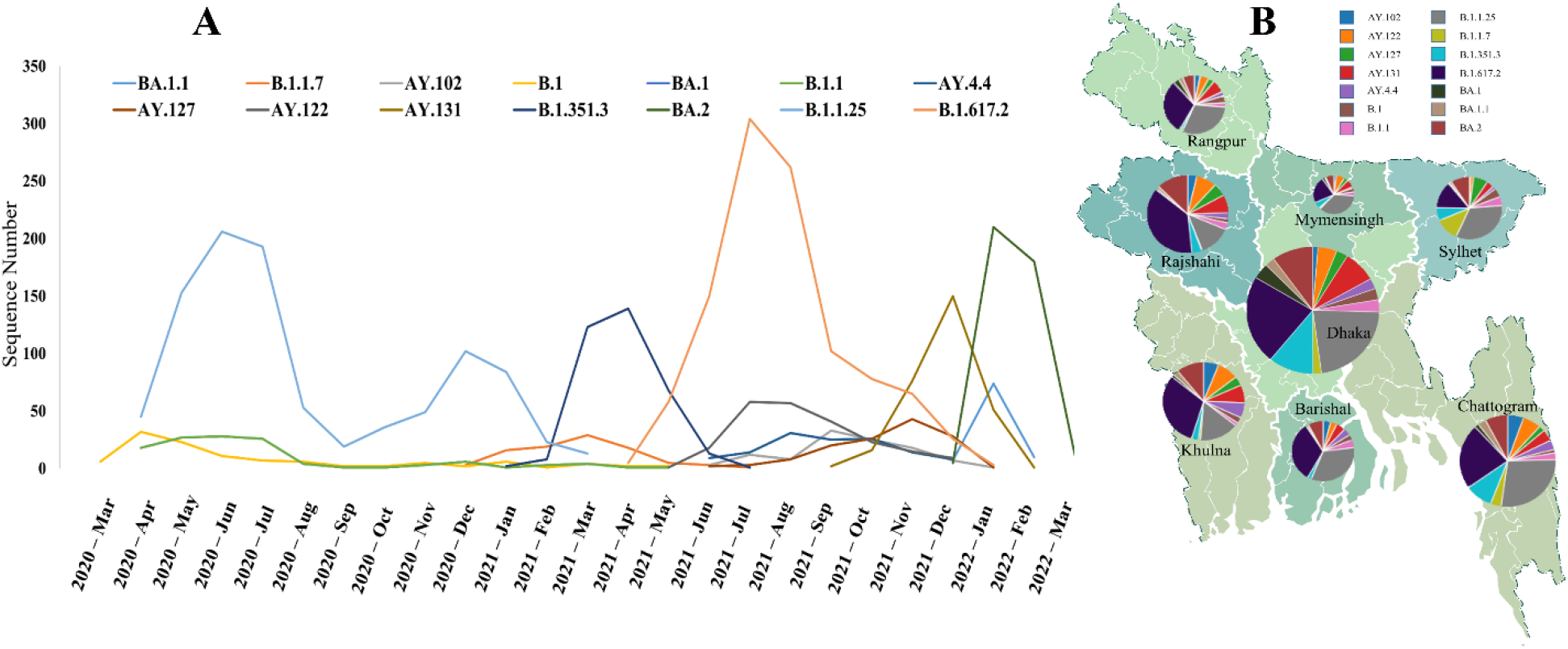
Distribution of major lineages in Bangladesh. **A**. Geographic distribution of major lineages at eight administrative divisions of Bangladesh. Dhaka contained the highest number of sequences and maximum diversity, while Mymensingh was the least diverse zone. **B**. Temporal distribution of major lineages. Maximum diversity was observed after the introduction of the Delta variant due to the emergence of different sub-lineage of it. Three major peaks depict the three variants responsible for three COVID-19 waves in Bangladesh.

According to our analysis, B.1.617.2 was the most dominant strain within a month after its introduction in April 2021. A number of distinct A lineages have also been observed, which were mostly sub-lineages of the delta variant, possibly due to the increased transmissibility of the variant. Specifically, AY.122 increased significantly from September 2021 while B.1.617.2 was declining. Meanwhile, the AY.131 lineage first appeared in Bangladesh in October 2021 and dominated all other variants in November 2021; more than half the sequences of December 2021 came from this lineage. This variant was eventually replaced by another highly transmissible variant called Omicron (B.1.1.529). Omicron first emerged in Bangladesh in December 2021 and took over within a month. Throughout Bangladesh, the Omicron variant has been dominant since January 2022.

### Regional distribution of different lineages

We then conducted a chronological lineages dynamics analysis in order to determine whether the variants were distributed evenly across Bangladesh’s administrative divisions. In terms of geographical distribution, Dhaka had the most diversified sequences from all thirty-five lineages, followed by Chattogram from 31. On the contrary, Mymensingh and Rangpur were less diverse areas with sequences from only 22 and 24 lineages, respectively, where most of the lineages represented only one or two sequences (Fig 1B). The Alpha variant was first detected in the Sylhet division and has since spread to the other five divisions except for Barishal and Rangpur, where the Delta and Omicron variant first appeared in Dhaka. Overall, the ratio of the dominating lineages was similar throughout the country, and our analyzed transmission network reflects that Dhaka was the hub of viral spread (Fig 2). Area-specific detailed chronological distribution of SARS-CoV-2 variants is provided in the supplementary file (Supplementary file 2).

**Fig 2:**
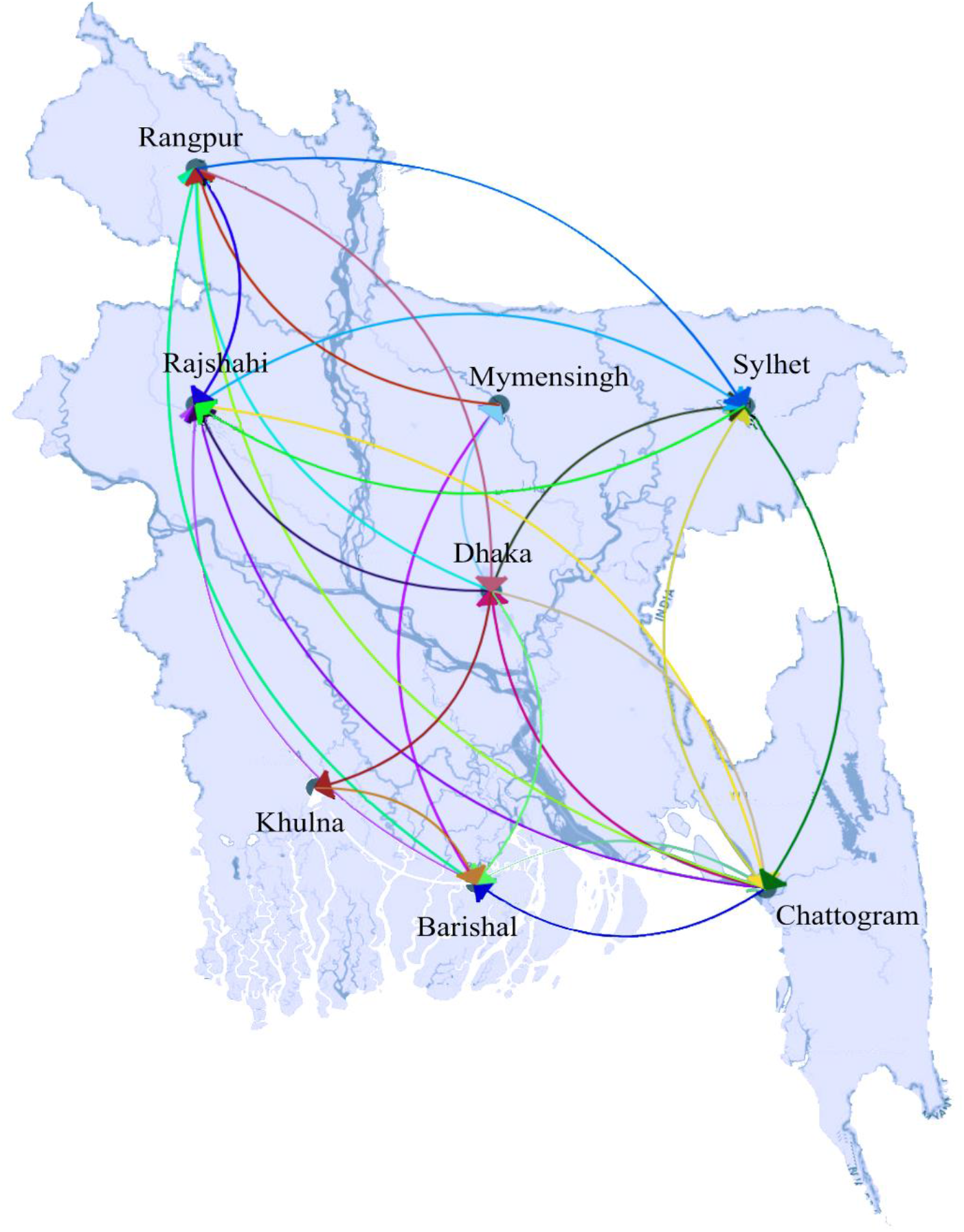
Transmission of SARS-CoV-2 in Bangladesh. Unlike others, Dhaka was connected with all other parts of the country, therefore recognized as a viral transmission hub. Arrows at the tip of the line dictate the direction of transmission.

To get a clearer idea of the viral circulation trend and back and forth transmission in different divisions of the country, we extensively analyzed the variants chronologically. We figured out that the whole country was mostly filled with a few major lineages throughout the times, but interestingly their dominance varied. We have seen that some lineages were missing from a particular area at a particular time and then returned, maybe due to mass people’s movement from other areas. For example, B.1.1 lineages were present in Mymensingh from the very beginning till June 2020. Then, this variant was missing there for five months but reappeared in the middle of December 2020. However, the variant was found during this period in Dhaka and Chattogram. On the other hand, the sub-lineages of Beta variant B.1.351.3 were missing in Sylhet for two months from February to March 2021 and occurred again in April 2021, while this variant was present in other divisions during this time. Several other back and forth circulation of strains were observed, for example, AY.100 and AY.102 in Dhaka. Detailed circulation of the variants information is provided in the supplementary file (Supplementary file 2).

Finally, we have built a viral transmission network using all our analysis data set sequences. Dhaka was found to be the center of viral transmission and directly connected with all other locations, while others were not. For example, we did not find any direct connection between Chattogram with Khulna and Mymensingh, Rangpur with Khulna, and Barisal did not have any connection with Sylhet (Fig 2). In addition, a strain-specific transmission network reveals the connections among different clusters and routes of viral spread from root to tip (Fig 3). With the time-calibrated analysis, we have observed that the sequences from Dhaka remain at the center of the network and determine the course of transmission forming connections with several subgroups.

**Fig 3:**
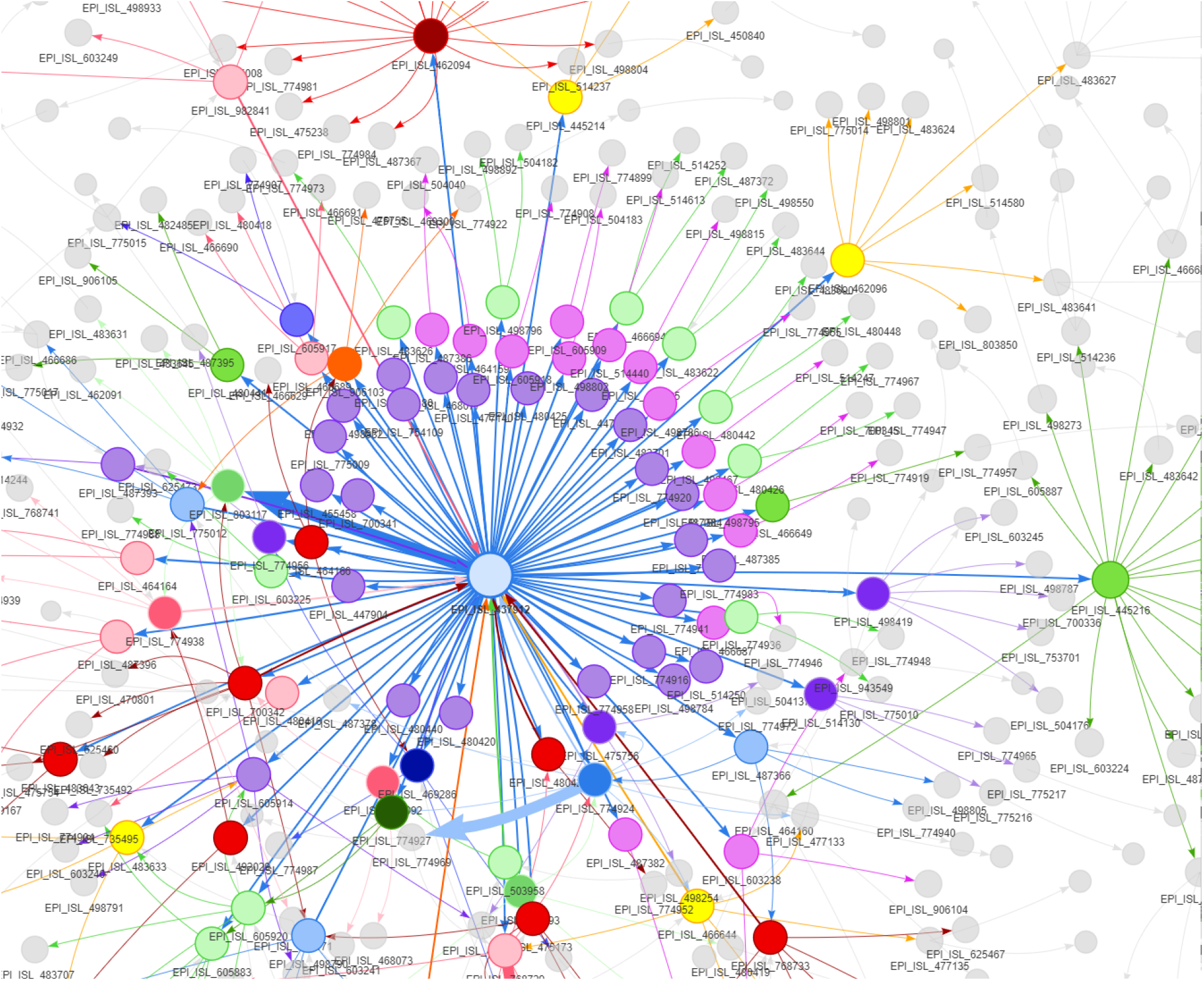
Strain to strain transmission network of the SARS-CoV-2 in Bangladesh. The sizes of nodes are proportional to the number of sequences a cluster contains and the thickness of the lines and arrows represent the frequency of transmission. The arrows reflect the direction of transmission among the viral clusters. The first sequence from the country is at the center of the network, and different clusters originated from the very first sequence, which gave rise to further subgroups; eventually, tips of the network reached.

### Mutation analysis summary

Till the present study, we have found 7659 unique mutations present in 4622 sequences where 482 were extragenic mutations, and the rest were in the coding regions. In the coding region, a total of 4103 missense, 2865 synonymous, ten insertion, 125 deletion and 74 premature stop codon mutations were observed (Fig 4A). Moreover, our analysis demonstrated 37.64 mutations per sequence, where 24.61 mutations were missense, and the ratio of acquiring missense over synonymous mutations increased gradually (Fig 4B). We have seen the number of mutations increased gradually over time, yet nearly 29% of the sequences carried mutations below 30, and more than 55.25% of sequences had 30 to 50 mutations. The highest number of mutations detected was 78 in two strains isolated from Dhaka on 28^th^ February 2022, and the lowest number was only one found in a sequence from 11^th^ May 2021. Fig 4B clearly demonstrates two remarkable rises in mutations, one in February 2021 due to the introduction of Delta variants. Another sharp rise was observed in January 2022 because of the highly transmissible Omicron variant with a large number of mutations in the spike protein. However, the individual genes went through mutation distinctively. Therefore, we thoroughly carried out the mutational analysis of all the SARS-CoV-2 sequences from Bangladesh and summarized the results in table 1 and figure 4.

**Fig 4:**
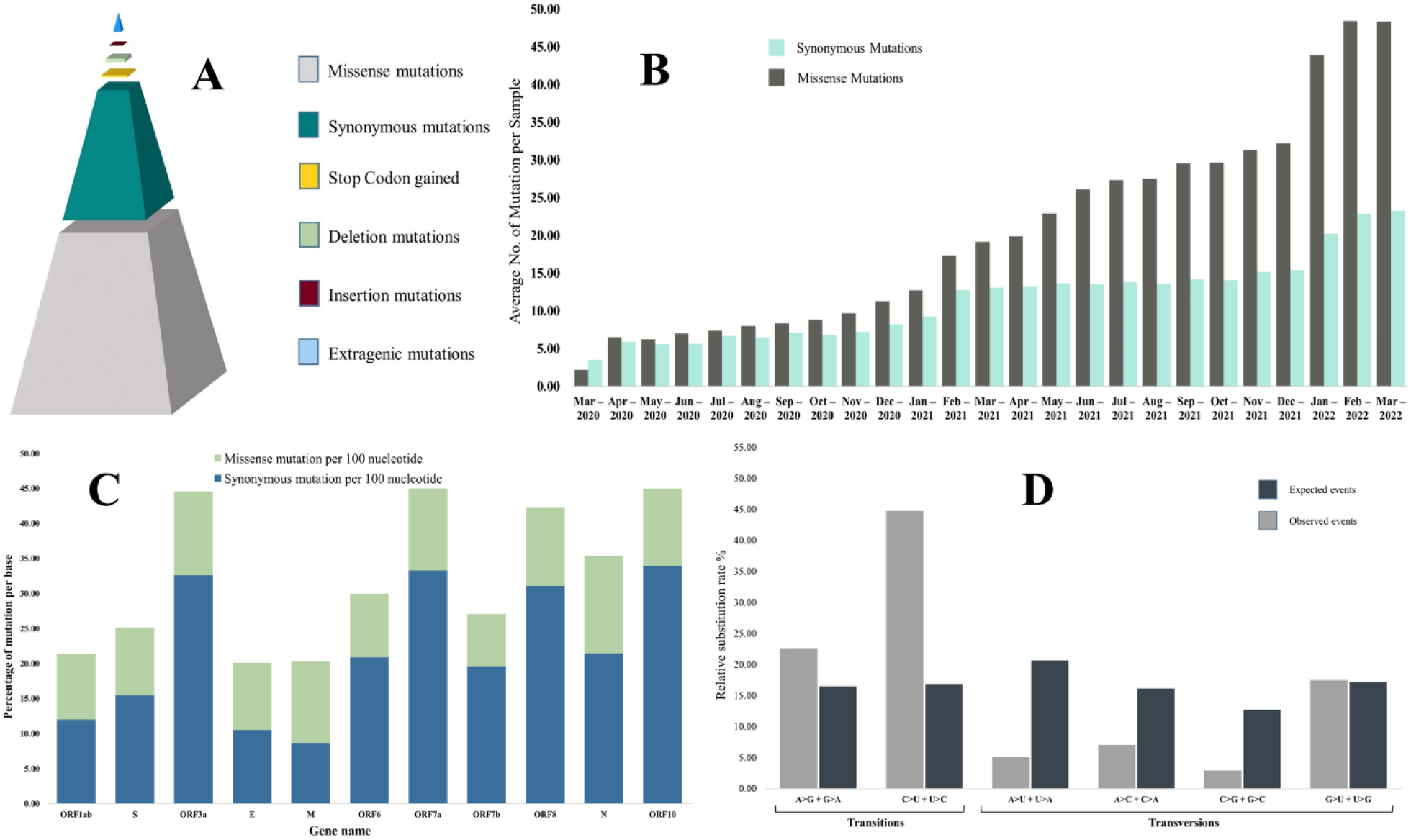
Summary of mutational events. **A**. Type of mutations among the sequences. Considering all the unique mutations, missense mutations were found to be the most prevalent event. **B**. The average number of mutations per sequence in each month. Although the number of mutations gradually increased with time, we observed a sharp increase from January 2021 when the Alpha variant entered the country. Moreover, more non-synonymous mutations emerged with time than synonymous mutations. **C**. Percentage of mutation per base in each gene. ORF3a had the highest density of mutations, while Envelop protein is the least mutated. Missense mutations are more prevalent than synonymous mutations. **D**. Nucleotide substitution rates for each of the four nucleotides among the SARS-CoV-2 genomes. Transition events were more prevalent than transversion events. C>U substitution rate was more than three times higher than the expected rate.

**Table 1:** SARS-CoV-2 mutation summary on individual genes.

| ORF | No. of non-mutant sequences | Percentage of mutated sequences | No. of synonymous mutations | No. of missense mutations | Percentage of missense mutations | Mutation density | No. of frequent mutations (n >= 10%) | No. of insertion mutation | No. of deletion mutation | No. of stop codon gained |
| --- | --- | --- | --- | --- | --- | --- | --- | --- | --- | --- |
| ORF1ab | 1 | 99.98% | 1997 | 2550 | 56.08% | 21.36% | 29 | 2 | 33 | 20 |
| S | 2 | 99.96% | 370 | 590 | 61.46% | 25.12% | 31 | 4 | 40 | 11 |
| ORF3a | 873 | 81.11% | 99 | 270 | 73.17% | 44.57% | 4 | 2 | 5 | 1 |
| E | 3395 | 26.55% | 22 | 24 | 52.17% | 20.18% | 1 | 0 | 0 | 1 |
| M | 1423 | 69.21% | 78 | 58 | 42.65% | 20.33% | 3 | 0 | 1 | 2 |
| ORF6 | 3838 | 16.96% | 17 | 39 | 69.64% | 30.11% | 1 | 0 | 4 | 3 |
| ORF7a | 2258 | 51.15% | 43 | 122 | 73.94% | 45.08% | 3 | 0 | 11 | 11 |
| ORF7b | 2395 | 48.18% | 10 | 26 | 72.22% | 27.27% | 2 | 0 | 4 | 4 |
| ORF8 | 1604 | 65.30% | 41 | 114 | 73.55% | 42.35% | 1 | 1 | 11 | 13 |
| N | 78 | 98.31% | 175 | 270 | 60.67% | 35.32% | 12 | 1 | 15 | 3 |
| ORF10 | 4250 | 8.05% | 13 | 40 | 75.47% | 45.30% | 0 | 0 | 1 | 5 |

ORF10 and ORF7a harbored the highest mutation density with 45.30% and 45.08% mutations per base, respectively, although only 8.05% of sequences were found to carry mutations in ORF10. On the other hand, 99.98% and 99.96% of sequences had mutations in ORF1ab and S genes, but their mutation density was lower at 21.36% and 25.21%, respectively. ORF6 was found to be the most stable gene of SARS-CoV-2 in sequences from Bangladesh, with only 16.96% sequences carrying the mutations, 30.11%% mutations per base and 69.64% missense mutations. ORF3a was identified to harbor the highest percentage (75.69%) of missense mutations. In comparison, the least percentage of missense mutations (42.65%) with 22.33% mutations per base was found in membrane protein-encoding gene M. It was clearly evident that non-structural proteins were subjected to more missense mutations than non-synonymous mutations compared with structural proteins (Fig 4C). In addition, we have found several deletions and insertion mutations where both the highest occurrences were found in the spike protein-coding S gene with 40 unique deletions and four insertions (Table 1). On the other hand, the highest number of unique stop codons were present in ORF1ab, with 40 out of 74 total stop codon mutations detected (table 1).

Among the 7786 mutations, 6968 were SNP, where 4697 and 2271 were involved in transition and transversion events, respectively, rendering a transition transversion ratio of 2.07. Transition mutations were calculated to be more prevalent than expected if mutational events took place randomly, which clearly revealed the nucleotide substitution bias (Fig 4D). Then, transition mutation C>U was the most frequent event, being 30.67% of total mutations and 45.50% of transition mutations, followed by the transversion event G>U, which was 15.37% of the total mutations (Supplementary file 3).

Then, out of the ten most prevalent mutations in Bangladesh, three were extragenic, one was synonymous, and six were missense mutations, where 23403A>G (missense mutation) was the highest prevalent, followed by the second highest 14408C>T (missense mutation) which resembles the global scenario and these two mutations were accompanied by 3037>C>T (synonymous mutation). Among the top 7 mutations in the coding region, three were in the spike protein (D614G, P681R and T478K), two were in the ORF1ab (P4715L and F924F), one was in membrane protein (I82T), and another was in ORF3a (S26L). These seven mutations had a high prevalence globally because these belong to different variants of concerns. In addition, 1163A>T (nsp2: I120F) was a highly prevalent and unique mutation found in Bangladesh from the beginning of the pandemic while it was absent in other countries. This mutation was present in more than 21% of sequences. Interestingly enough, from linkage disequilibrium (LD) analysis, we have found that all the mentioned mutations had a very strong correlation (R^2^ =1.00) and occurred in parallel since their first appearance (Fig 5). 1163A>T mutation was predicted to have been 100% (R^2^ =1.00) connected with 20 more mutations in parallel, considering the high frequent mutations present at least in 10% of our sequence set. Additionally, several other mutations were occurring in parallel with very strong LD values due to the introduction of several variants of concerns in the country (Fig 5).

**Fig 5:**
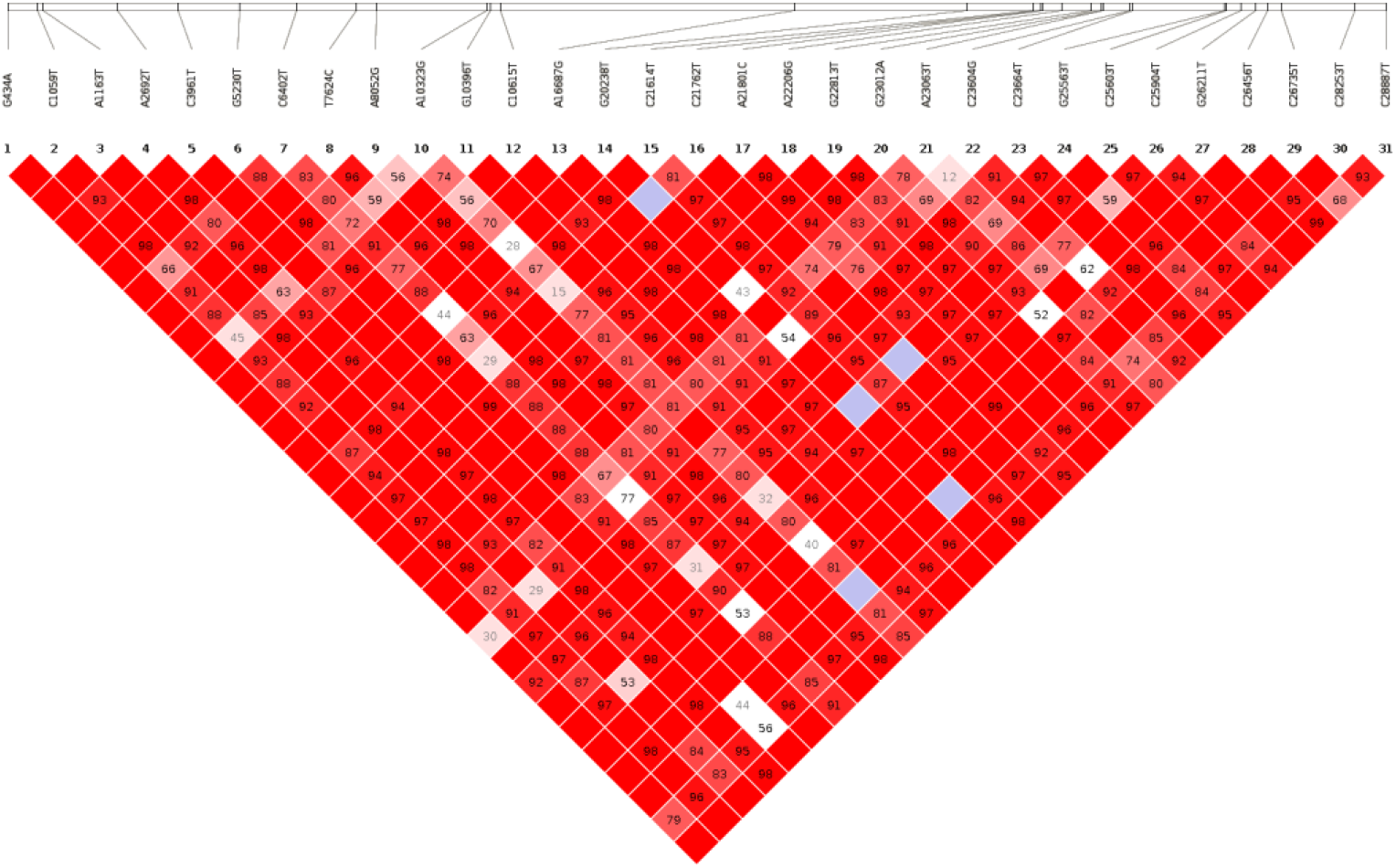
Linkage disequilibrium plot. The LD plot is generated considering the most prevalent SNPs. The number at the top denotes the SNP position, and squares are colored by standard (D’/LOD). The brighter red color indicates a higher D′ value and vice versa. The number in square is r ^2^ value.

### Effect of the mutations

The mutations affect viral infectivity, transmissibility, virulence, viral fitness, selection pressure, proteome structure, and evolution. Overall, the SARS-CoV-2 genomes had very high nucleotide identity with an average of 37.64 mutations and low overall nucleotide diversity (π) 0.004. Although overall nucleotide diversity was lower, it varied from gene to gene. For example, ORF8 had the highest nucleotide diversity (0.01543), while gene ORF10 was most stable with a π value of 0.00059 (Table 2). Analyzing the Bangladeshi sequences, the most diverse spot of the genome was in the spike protein gene at position 23009 with nucleotide diversity value π=0.16372 while the least diverse spot found was at position 11069 of ORF1ab with a π value of 0.00005. Nevertheless, most of the genes overall had lower nucleotide diversity, which signifies selective sweep due to increased mutations that benefit the strains and lead to the reduction of genetic variation. In addition, we have found that three genes (ORF3a, ORF7b and ORF10) were under positive selection pressure or directional selection because the mutations present in them were advantageous to them; therefore, their frequencies were on the rise while the rest of the genes were facing negative evolution pressure to stabilize against the deleterious mutations they have got from random mutational events. Precisely, only 47 sites were facing positive or diversifying selection pressure against 190 sites found to be under negative or purifying pressure to stabilize the genomic variations (Supplementary file 3).

**Table 2:**
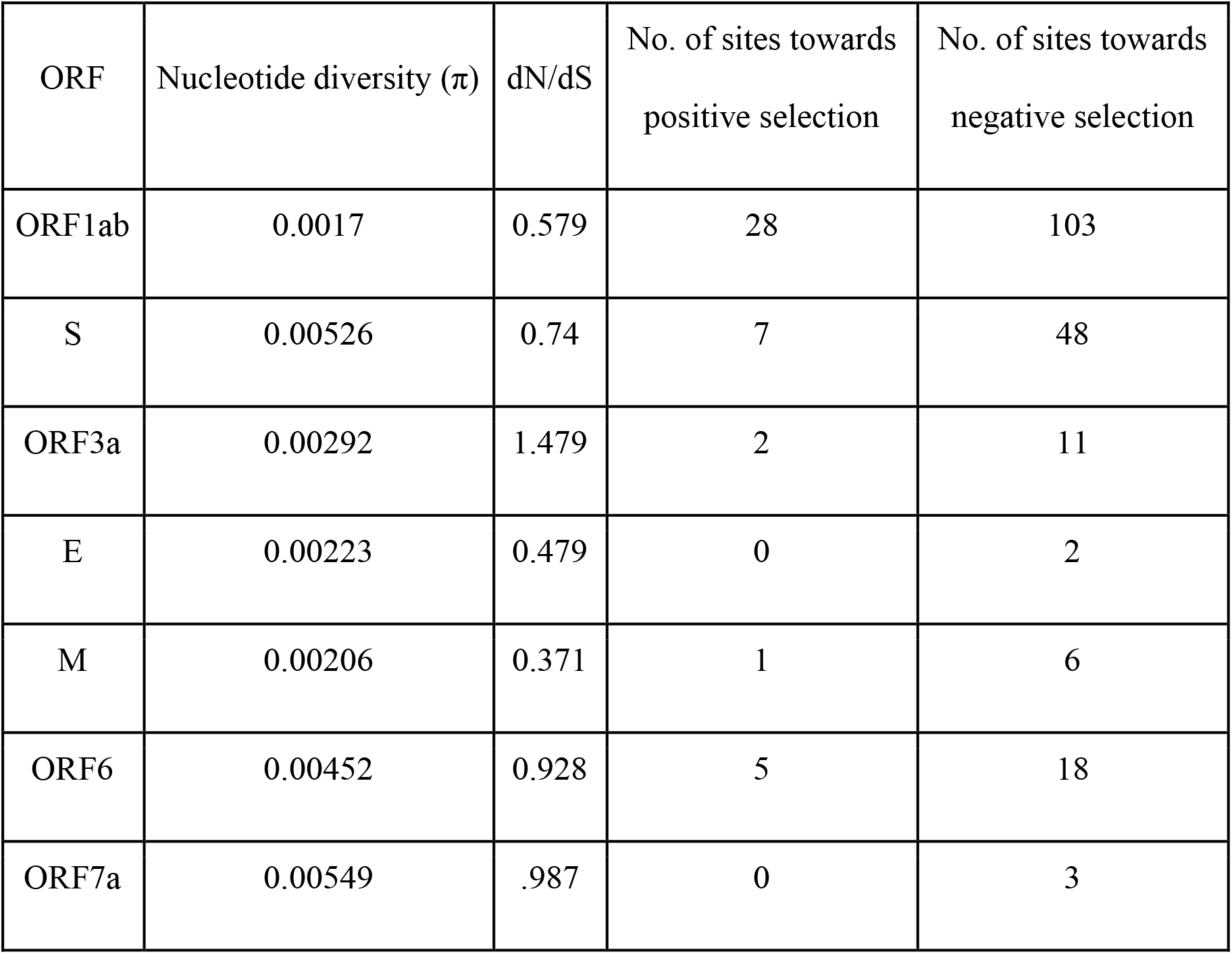
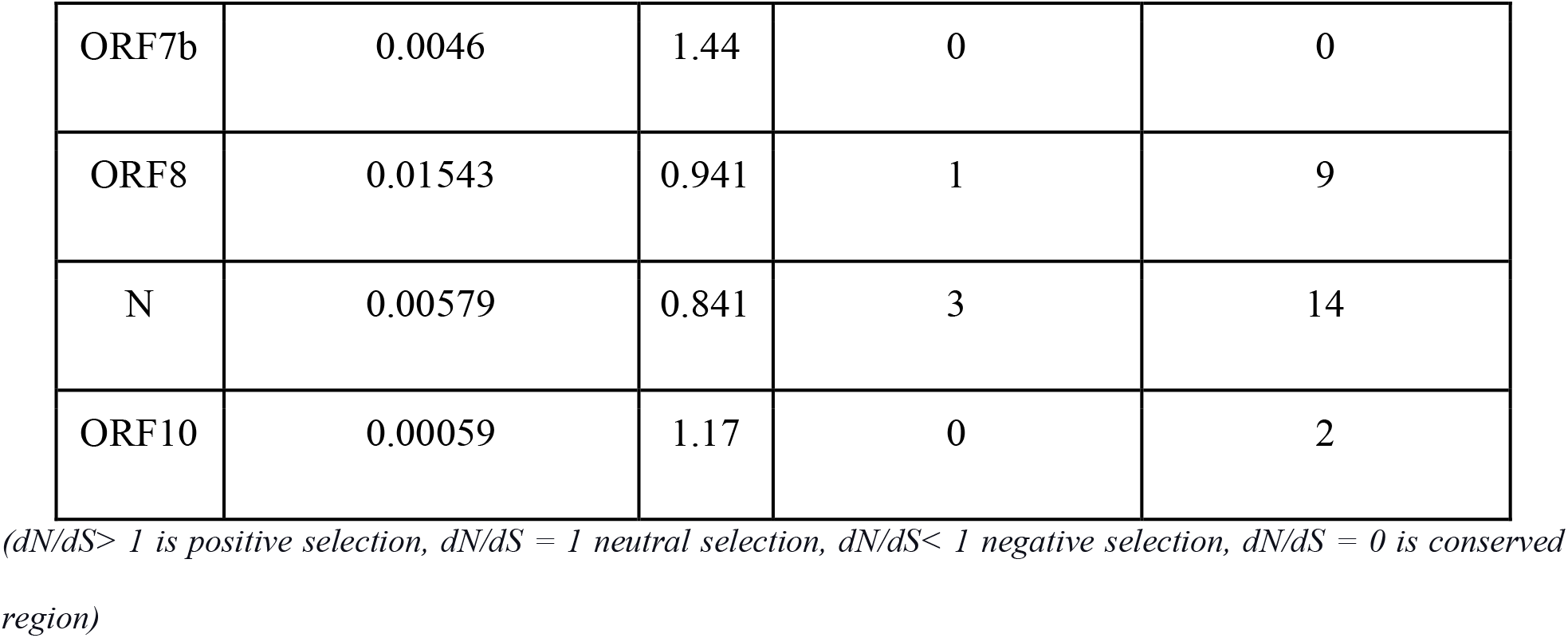
Summary of Mutation’s effect on each protein.

| ORF | Nucleotide diversity ($\pi$) | dN/dS | No. of sites towards positive selection | No. of sites towards negative selection |
| --- | --- | --- | --- | --- |
| ORF1ab | 0.0017 | 0.579 | 28 | 103 |
| S | 0.00526 | 0.74 | 7 | 48 |
| ORF3a | 0.00292 | 1.479 | 2 | 11 |
| E | 0.00223 | 0.479 | 0 | 2 |
| M | 0.00206 | 0.371 | 1 | 6 |
| ORF6 | 0.00452 | 0.928 | 5 | 18 |
| ORF7a | 0.00549 | .987 | 0 | 3 |
| ORF7b | 0.0046 | 1.44 | 0 | 0 |
| ORF8 | 0.01543 | 0.941 | 1 | 9 |
| N | 0.00579 | 0.841 | 3 | 14 |
| ORF10 | 0.00059 | 1.17 | 0 | 2 |

( $dN/dS > 1$ is positive selection, $dN/dS = 1$ neutral selection, $dN/dS < 1$ negative selection, $dN/dS = 0$ is conserved region)

Finally, these mutations affected the virus from the evolutionary perspective and shook the stability of the proteins they encode. Most of the mutations were previously reported to affect the stability of the whole proteome of SARS-CoV-2 negatively. However, all the genes were not affected to the same extent by mutational events (Fig 6). For example, only 42.65% of mutations on the membrane protein-coding M gene were missense which was 73.55% in the case of ORF3a (table 1). As of now, vaccines and therapies target the spike protein, which is highly mutated. That’s one reason why people continue to develop symptoms after successful vaccination. It is possible that current vaccines and therapies will not work in future due to a high number of mutations occurring. The less affected genes could therefore be targeted for medicine and vaccine development. Figure 6 shows spikes that represent mutations, and the height of the spikes is proportional to the number of mutations that have taken place at that position in the genome. As we can see, there are plenty of stable regions between the spikes, which could be targeted for therapeutics and vaccine development against SARS-CoV-2.

**Fig 6.**
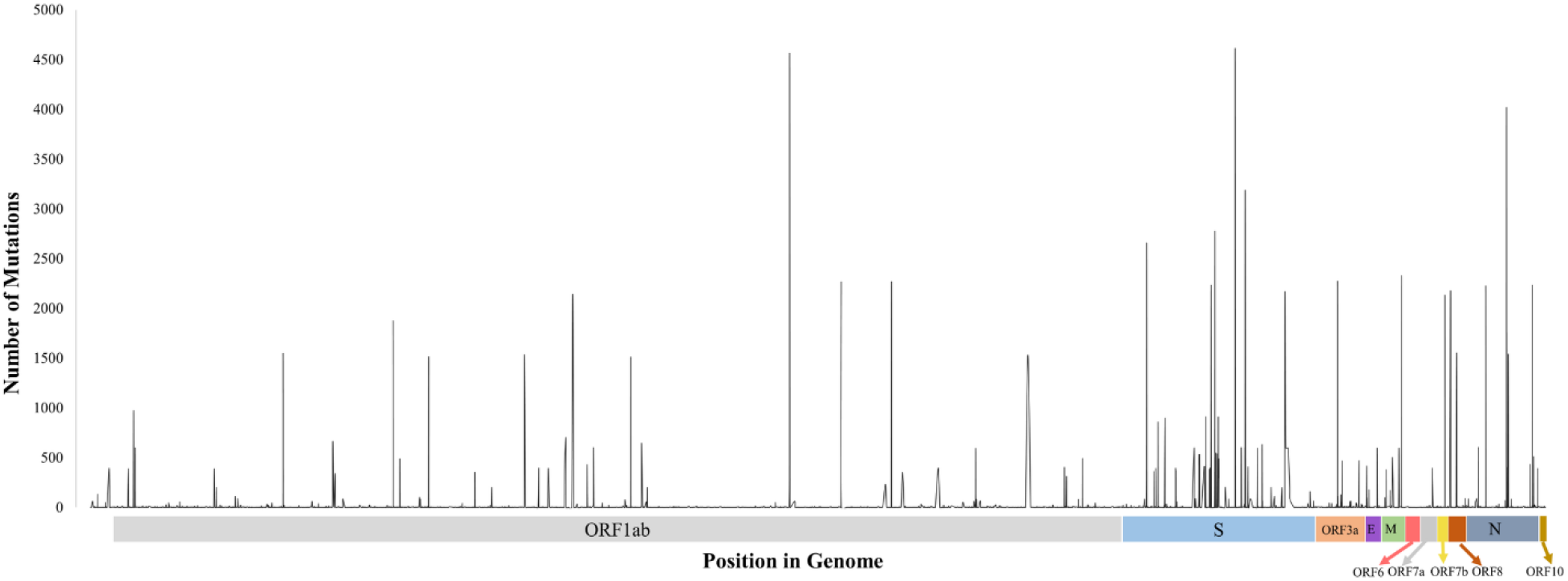
Distribution of missense mutations in the genome. The height of the spikes is proportional to the number of sequences that got mutation at that location. Regions between these spikes are stable, which could be targeted for further vaccine and therapeutics development.

## Discussion

SARS-CoV-2 has been circulating in Bangladesh for over two years, and many strains are sequenced from different parts of the country, helping us carry out downstream analysis to depict different variants, transmission, and evolution inside the country. Investigating 4622 whole-genome sequences from Bangladesh, we have seen B.1.1.25 lineage was in dominance since the beginning along with other lineages containing a small number of sequences, but since March 2021, strains from B.1.1.25 lineage are surmounted by another lineage B.1.351.3, which is a sub-line of Beta variant. In addition to that, we have observed a slight increase of Alpha variants for a short time since their first appearance in January 2021. However, In April 2021 Delta variant emerged and dominated other variants until the arrival of Omicron. The Omicron variant comprises three main sub-lineages termed BA.1, BA.2 and BA.3. Both BA.1 and BA.2 are found in Bangladesh, and currently, BA.2 is the dominant variant. Although BA.1 and BA.2 have numerous mutations in common, 20 mutations in the spike protein differentiate the two sub-lineages, and BA.2 also displays a marked decreased sensitivity to many neutralizing monoclonal antibodies (mAbs) when compared to previous VOCs (37). Therefore, with further mutations, this BA.2 sub-lineage is keeping the risk of having another COVID-19 wave alive in the country.

On the other hand, geographical analysis depicts Dhaka and Chattogram containing a more diversified number of sequences than other parts of the country. Our analysis has limitations at this point because we had a higher number of sequences from these two regions than others. The sequences were more diversified in the first phase of the pandemic. However, with the arrival of the Delta and Omicron variants, the divergence reduced drastically, maybe due to viral adaptation following the “Survival of the fittest’’ theory of natural selection, although we have found several sub-lineages of the Delta variant. Additionally, we have seen Dhaka being the viral transmission hub, which is obvious since it is the capital city of Bangladesh, but this city is not the only transmission source. From extensive analysis, we have built the SARS-CoV-2 transmission network between different administrative divisions and observed the back and forth transmission of the virus inside Bangladesh. This situation arose due to a lack of restriction on the mass movement; public gatherings were not limited duly, and other socioeconomic events.

From the mutational perspective, we have seen a total of 7659 unique mutations present in 4622 sequences with 37.64 mutations per sample where on average 24.61were coding variants, which happens to be significantly higher than the global average of 7.23, reported in July 2020 (38). This sharp rise of mutations indicates the SARS-CoV-2 might be facing strong challenges from the host’s immunologic response in addition to random regular mutational events of RNA viruses, which is one of the reasons for the emergence of new variants of concerns. At the nucleotide level, 67.41% of the mutations were transition events, and a molecular bias was present for C>U, which tends to mutate hydrophilic amino acids into hydrophobic ones (36). Overall, 1109 unique C>U missense mutation on the genome was observed, where 194, 289, and 162 of the mutations were skewed towards phenylalanine, isoleucine, and leucine codons, respectively. Moreover, 236 C>U mutations were involved in altering the proline, which is known to be a strong helix breaker (39). Therefore, proline to another amino acid shift might have a deleterious effect on the SARS-CoV-2 proteome.

On the other hand, we have found that most of the genes were under negative selection pressure while only three non-structural protein-coding genes were under Darwinian (positive) selection, which indicates that most of the random mutational events were deleterious for the SARS-CoV-2 (40), maybe due to the immunologic potential of people of Bangladesh and our demography. However, the dN/dS ratio of the receptor-binding region (RBD) of the spike protein was higher, indicating that mutations in this region were advantageous. The result correlates with the emergence of different variants of concerns like Alpha, Beta, Delta and Omicron. The RBD region is considered the most important part of the virus since it attaches to ACE2 during viral infection to host cells. Those advantageous mutations might increase pathogenicity, infectivity, transmissibility, and letting it evade the host immunity (41–43). Moreover, most of the current therapies and vaccines are developed targeting the BRD-ACE2 interaction. Therefore, a higher dN/dS ratio also warns us about the emergence of new deadly variants in future with further mutations in this region and vaccine failure. However, analyzing all the sequences from Bangladesh, we have seen that the whole proteome was not affected to the same extent. There were regions of high and low nucleotide diversity. While highly affected regions are evolving faster, regions with low nucleotide divergence would render us the opportunity to develop new vaccines and antibodies for treatment and development detection kits to reduce the number of false test reports during this pandemic.

To sum up, considering the limitations regarding sequence number variations in different parts of the country, we have analyzed all the unique and global mutations present in Bangladesh, which are thoroughly reported in the supplementary files. This data would facilitate researchers further from various perspectives like investigating viral transmission, the connection among isolates, evolution patterns, and dynamics of divergence of the virus.

## Methods and materials

### Sequence retrieval and Lineage determination

Using completeness and coverage filters on the sequences, all the SARS-CoV-2 genomes submitted from Bangladesh until 25th March 2022 were retrieved from the Global Initiative on Sharing All Influenza Data (GISAID) database (www.gisaid.org) (9). Prior to downstream analysis, all sequences were quality checked and sequences with more than 5% ambiguous characters were omitted. The sorted sequences were then classified by Phylogenetic Assignment of Named Global Outbreak LINeages (Pangolin) with COVID-19 Lineage Assigner (https://pangolin.cog-uk.io/) (20). Furthermore, we excluded lineages carrying less than ten sequences to address the important lineages. We analyzed and visualized the sequence lineage distribution in R within the country.

### Transmission analysis

First, the selected sequences were aligned using the Mafft algorithm (21), followed by the construction of a maximum likelihood phylogenetic tree using IQ-TREE (22)and calibrating the tree based on time with TreeTime (23). Using the StrainHub tool (24), we built the SARS-CoV-2 transmission network in Bangladesh from the reconstructed tree and metadata.

### Mutation Analysis

We have aligned each sequence with the reference sequence (NC_045512.2) (8) using the minimap2 algorithm (25) and called the variants with Samtools (26). Additionally, SNP-sites (27), CovSeq (28) and an online server Coronapp (29) were used to detect the mutations present in the sequences and the common mutations from these four sources were considered. Finally, SNPeff was used to predict the impact of the mutations (30).

### Effects of mutation

First of all, we used TASSEL software (31) to determine the nucleotide diversity (π) using a 20 base-pair window at five base-pair steps. Then we calculated the direction of selection in the sequences to know if diversity moves away from neutrality and to understand the pattern of evolution using the SLAC algorithm (32) in the HyPhy software package (33). Linkage disequilibrium among mutations prevalent in 10% or more sequences were calculated using AutoVem (34) and presented by the R2 index using HaploView (35). Then, along with determining the nucleotide substitution bias, the expected and observed transition, transversion events as well as their ratio were calculated by the method used by Matyášek R, Kovařík A (36).

## Acknowledgements

The authors are grateful for the efforts of all research institutes in Bangladesh in continuously sequencing the SARS-CoV-2 virus to track its changes and provide access to the data. We also would like to express our gratitude to all scientists, researchers, and health care workers around the world for their valuable contribution to the fight against this pandemic.

